# Longitudinal Morphological and Functional Characterization of Human Heart Organoids Using Optical Coherence Tomography

**DOI:** 10.1101/2022.01.19.476972

**Authors:** Yixuan Ming, Senyue Hao, Zhiyao Xu, Anna Goestenkors, Yonatan R. Lewis-Israeli, Brett D. Volmert, Aitor Aguirre, Chao Zhou

## Abstract

Organoids play an increasingly important role as *in vitro* models for studying organ development, disease mechanisms, and drug discovery. Organoids are self-organizing, organ-like three-dimensional (3D) cell cultures developing organ-specific cell types and functions. Recently, three groups independently developed self-assembling human heart organoids (hHOs) from human pluripotent stem cells (hPSCs). In this study, we utilized a customized spectral-domain optical coherence tomography (SD-OCT) system to characterize the growth of hHOs. Development of chamber structures and beating patterns of the hHOs were observed via OCT and calcium imaging. We demonstrated the capability of OCT to produce 3D images in a fast, label-free, and non-destructive manner. The hHOs formed cavities of various sizes, and complex interconnections were observed as early as on day 4 of differentiation. The hHOs models and the OCT imaging system showed promising insights as an *in vitro* platform for investigating heart development and disease mechanisms.

## 1. Introduction

In recent years, organoids have gained increasing attention as *in vitro* three-dimensional (3D) models for investigating organ development, disease mechanisms, regenerative medicine, drug toxicity, and many other applications (Abbott 2003; Friedrich et al. 2009; Sutherland 1988; Takebe and Wells 2019; Weiswald et al. 2015). Organoids are self-organizing *in vitro* models that consist of organ-specific types of cells, mimic organ functions, and form organ-like structures (Clevers 2016). Organoid models have been developed for many human organs, such as the brain (Mansour et al. 2018; Quadrato et al. 2016; Wang 2018), liver (Artegiani et al. 2019; Vyas et al. 2018; Wang et al. 2018), kidney (Garreta et al. 2019; Homan et al. 2019; Takasato et al. 2016), intestine (Finkbeiner et al. 2015; Kasendra et al. 2018; Petersen et al. 2014), lung (Dye et al. 2015; Miller et al. 2019), placenta (Deloria et al. 2020), retina (Capowski et al. 2019) and heart (Drakhlis et al. 2021; Hofbauer et al. 2021; Keung et al. 2019; Kupfer et al. 2020; Lewis-Israeli et al. 2021b; Li et al. 2018; Mills et al. 2017). Organoids recapitulate critical *in vivo* features, including the diffusion of nutrients, oxygen and metabolic waste, the cell microenvironment, and interactions among cells, and they have been proven to promote gene expression, develop organ-specific types of cells, mimic organ functions, and form organ-like structures (Clevers 2016). In the development of organoid models for human organs, human heart organoids (hHOs) have gained less progress due to the structural complexity of the human heart (Lewis-Israeli et al. 2021a). Early studies of hHOs relied on 3D scaffolds, on which human stem cells or differentiated cardiac cells were cultured and guided to form pre-designed 3D structures (*e*.*g*., direct assembly). Functional heart chambers were reported in hHOs fabricated in this way (Keung et al. 2019; Kupfer et al. 2020; Li et al. 2018; Mills et al. 2017). However, heart organoids fabricated via direct assembly often fail to capture important *in vivo* features, including multiple cell types, the cell ratio and density, complicated morphology, and physiological development (Lewis-Israeli et al. 2021a). Recently, self-assembling hHOs have been reported (Drakhlis et al. 2021; Hofbauer et al. 2021; Lewis-Israeli et al. 2021b). In these hHOs, stem cells form spherical 3D structures via self-organization. *In vitro* differentiation and development have been achieved through exposure to small molecules and growth factors. Self-assembling hHOs have generated diverse cell types, gene expression profiles associated with *in vivo* developmental stages, and complicated morphology, such as chambers, vascularization, layered structures, and atrioventricular specification (Drakhlis et al. 2021; Hofbauer et al. 2021; Lewis-Israeli et al. 2021b). Compared with direct-assembly methods, self-assembly methods have allowed for diversity of the organoid models, but also have resulted in variation in the overall morphology and cell ratio, and less control of the maturation state of the models.

Morphogenesis of hHOs is of great interest, as organoids are expected to capture the development of heart-specific structures, including functional chambers, blood vessels, heart valves, and so on. Morphogenesis of the organoids can be studied using fluorescence imaging, where organoids are fixed and labeled with fluorescent markers for specific cell types. Self-assembling hHOs are reported to have densely packed cells and sizes ranging from several hundred micrometers to one millimeter in diameter (Drakhlis et al. 2021; Hofbauer et al. 2021; Lewis-Israeli et al. 2021b). Limited by the penetration depth of both fluorescent markers and the imaging light, only a small portion of morphological information can be retrieved if the organoid is processed and imaged as a whole. Tissue sectioning can address this difficulty, but it requires extra labor for sample preparation, image acquisition, and data processing, and it loses 3D information. Besides, both immunostaining and tissue sectioning terminate the organoids, making it impossible to track the same organoid throughout all developmental stages. Thus, a fast and non-invasive imaging modality is needed to better characterize the morphology of hHOs.

Optical coherence tomography (OCT) is a fast, non-destructive, and label-free imaging modality that has been demonstrated in studying various organoid models (Capowski et al. 2019; Deloria et al. 2020; Huang et al. 2017; Lewis-Israeli et al. 2021b). OCT detects intrinsic backscattered light signals and yields high contrast signals at interfaces with different materials. Thus, the morphology of the organoids, such as structural surface variations, cavity formations, vascularization, and layers of different cell types can all be visualized via OCT. OCT studies of trophoblast organoids derived from human placentas revealed that cavities within the organoids was associated with syncytiotrophoblast formation (Deloria et al. 2020). OCT observations of structural hallmarks in human retinal organoids, including a bowl-like contour and hollow cores, indicated retinal organoids at different developmental stages (Capowski et al. 2019). OCT successfully detected necrotic regions within tumor organoids based on the intensity attenuation profile of their signals (Huang et al. 2017). Besides 3D morphological information, OCT can be configured to detect variations of the same cross section (B-scan) versus time at high frame rates, which makes OCT a promising tool for tracking beating patterns of hHOs. In this study, OCT imaging, together with a newly reported protocol to generate self-assembling hHOs (Lewis-Israeli et al. 2021b), is demonstrated as a promising *in vitro* platform to investigate morphogenesis and 3D beating patterns, among other features of hHOs.

## 2. Materials and methods

### 1.1. Stem cell culture

The stem cell line used in this study was a wild-type C human induced pluripotent stem cell (WTC-hiPSC) line with genetically engineered GCaMP6f (Chen et al. 2013; Huebsch et al. 2015). hiPSCs were cultured in 6-well plates coated with growth factor reduced Matrigel (Corning) and maintained in Essential 8 Flex medium (E8, Gibco) at 37 °C in a humidified incubator supplied with 5% CO_2_. After reaching 60-80% confluency, hiPSC cultures were passaged into new 6-well plates or collected to fabricate hHOs. After at least three passages post-thawing, hiPSC cultures were ready for hHOs fabrication.

### 1.2. Human heart organoids (hHOs) fabrication

A detailed fabrication protocol was described in our previous work (Lewis-Israeli et al. 2021b). A schematic of the protocol is shown in Figure 1A. Briefly, hiPSCs were dissociated into single cells with Accutase (Innovative Cell Technologies), resuspended in E8 Flex medium with 2 μM Thiazovivin (Tocris) at 100,000 cells/mL, and seeded at a volume of 100 μL per well in 96-well ultra-low attachment microplates (Corning). After centrifuging at 100 g for 3 min, microplates were placed into a humidified incubator maintained at 37 °C and 5% CO2 (day -2). After 24 hours, 50 μL of the medium was replaced with 200 μL of fresh E8 Flex medium (day -1). 24 hours later (day 0), 66% of the medium (or 166 μL) was replaced with RPMI 1640 medium (Gibco) containing 2% B-27 supplement, minus insulin (Gibco), 4 μM CHIR99021 (Selleck), 0.36 pM Recombinant Human Bone Morphogenetic Protein 4 (BMP4, Gibco), and 0.08 pM Activin A (Fisher Scientific). After another 24 hours (day 1), 166 μL of the medium was replaced with fresh RPMI 1640 medium with B-27 (RPMI 1640/B27), minus insulin. On day 2, 166 μL medium was removed, and the same volume of fresh RPMI 1640/B27, minus insulin, and 2 μM Wnt-C59 (Selleck) was added. Organoids were incubated for 48 hours, and then 166 μL of the medium was refreshed with RPMI/B27, minus insulin (day 4). On day 6, 166 μL of media was replaced with RPMI/B27 (with insulin). On day 7, a one-hour exposure of 2 μM CHIR99021 was performed. After day 7, the culture medium was changed every 48 hours with fresh RPMI/B27. During medium change, only half or 66% of the medium was removed from each well to minimize disturbance and stress generated during the process.

**Figure 1.**
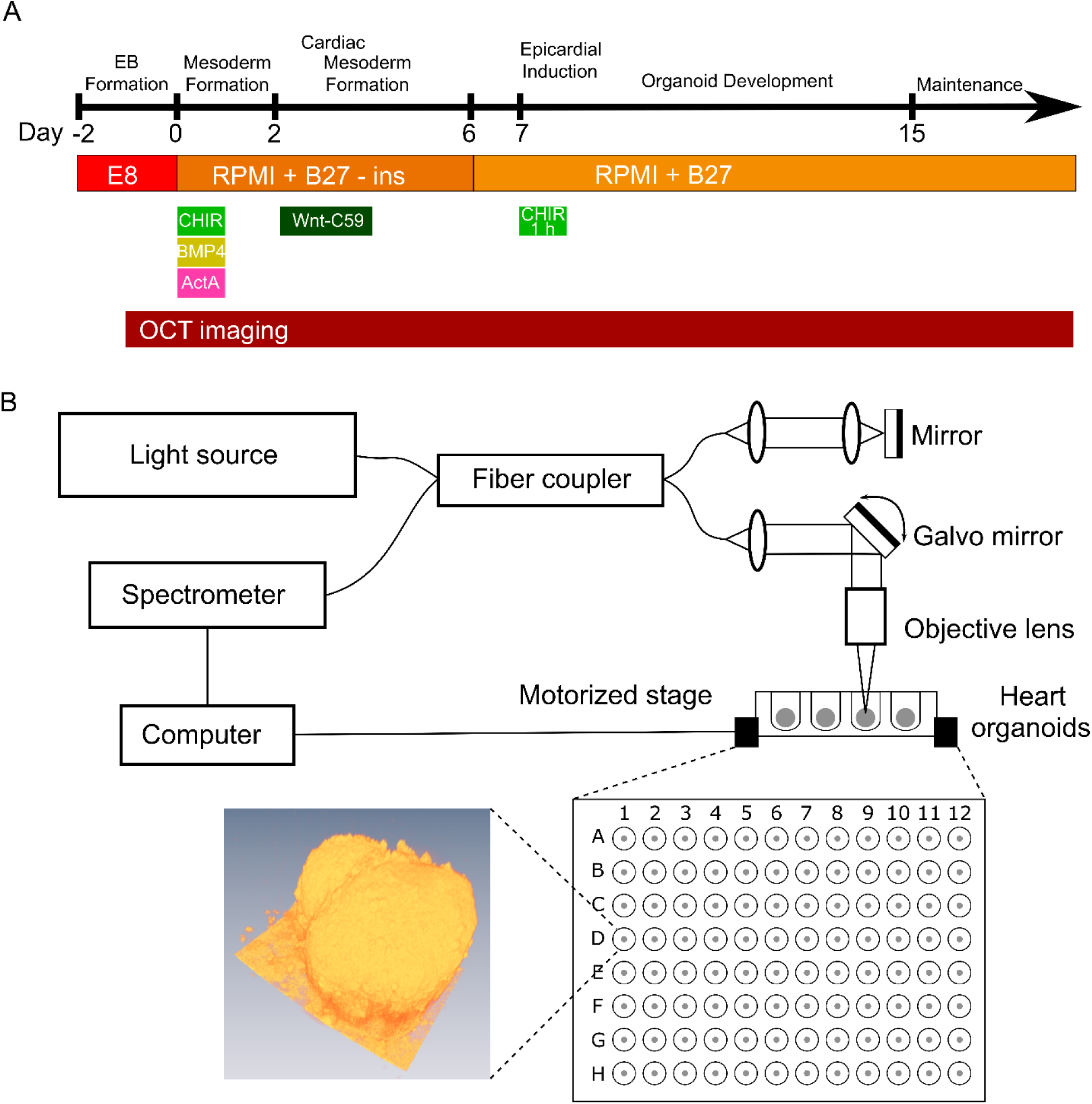
Experimental setups. (A) Schematic of the protocol for human heart organoid fabrication. OCT was performed every day from day -1 to day 22, and every other day from day 24 to day 30. (B) Schematic of the OCT setup. Bottom left shows a 3D rendering of an hHO on day 27. Abbreviations: E8, Essential 8 Flex medium; RPMI, RPMI 1640 medium; CHIR, CHIR99021; BMP4, Recombinant Human Bone Morphogenetic Protein 4; and ActA, Activin A (see Materials and methods for details).

### 1.3. Optical coherence tomography (OCT)

The detailed OCT setup can be found in our previous work (Huang et al. 2017). A schematic of the OCT setup is shown in Figure 1B. In this customized spectral-domain OCT (SD-OCT) setup, a superluminescent diode (Thorlabs, SLD1325) with a 1320-nm central wavelength and 110-nm spectral range was used as a broadband light source. Signals were detected by a spectrometer, which contained a 1024-pixel InGaAs line-scan camera (Sensors Unlimited, SU1024LDH2) with up to 20 kHz A-scan rate. The transverse resolutions were estimated as ∼7 μm using an USAF resolution test target (Thorlabs). Axial resolution was estimated as ∼4.9 μm, assuming the refractive index of heart tissue as 1.38 (Dirckx et al. 2005). Samples were imaged with a 5X objective lens. A motorized sample stage was controlled by the computer and held a 96-well plate in which heart organoids grew.

To further improve the resolution of the OCT, the light source was replaced with a supercontinuum laser (Fianium SC450) with a wavelength range from 400-2000 nm. For OCT imaging, we used a wavelength range from 1175 nm to 1420 nm, centered at 1300 nm. The spectrometer was upgraded to a Cobra 1300 (Wasatch Photonics), which contains a 2048-pixel InGaAs line-scan camera (Sensors Unlimited, GL2048) with up to a 147 kHz A-scan rate. Transverse and axial resolutions were improved to ∼3.48 μm and ∼2.36 μm, respectively. To track the beating of human heart organoids, the slower scanning direction of the galvo system was grounded so that OCT scanned only the same cross-section (B-scan) over time. Orthogonal B-scanning was performed by switching the two control signals to the galvo system. The number of A-scans was set as 600 per B-scan, and the exposure time was set as 94 μs. The equivalent frame rate was ∼15 frames per second (fps).

Starting from day 5 of the hHOs’ development, the OCT exposure time was set as 1 ms to improve the imaging depth. Acquired OCT images were first loaded in ImageJ (Schneider et al. 2012) for scale correction and then in Amira (Thermo Fisher Scientific) for segmentation and 3D rendering. Segmentations of the whole organoids were used to quantify the volumes of the organoids and cavities. The voxels in each frame were summed to obtain the volumes in voxels, and the final volumes were calculated by multiplying the volume of the voxel (mm^3^/voxel). Volumes of the organoids were calculated by adding the volumes of cellular materials and cavities.

### 1.4. Calcium imaging

To perform calcium imaging, wild-type C human induced pluripotent stem cells (WTC hiPSCs), expressing calcium indicator GCaMP6f (Chen et al. 2013; Huebsch et al. 2015), were used to fabricate hHOs. An inverted fluorescence microscope (IX71, Olympus) was used to record calcium transients of hHOs at around 10 fps. Acquired data were processed with customized MATLAB programs. Briefly, 10-by-10-pixel binning was first applied to the fluorescence recordings to reduce motion artifacts introduced by beating of the hHOs. Next, the fluorescence intensity *F* of each bin was calculated as the mean grey value of the bin. The baseline *F*_0_ of the tracing was calculated using asymmetric least squares smoothing, published in a previous work (Eilers and Boelens 2005). The fluorescence change Δ*F*/*F*_0_ was calculated using Δ*F*/*F*_0_ = (*F* − *F*_0_)/ *F*_0_.

A customized MATLAB program was used to analyze the propagation of calcium signals. First, the region of interest (ROI) was drawn to cover only the hHOs from the corresponding bright field recording. A threshold of the calcium signal intensity was set so that pixels outside the ROI were excluded from analysis. The time instant of the initiation of calcium transients for each pixel within the heart organoid was recorded. Within the same beating cycle, the initiation times of the calcium transients for each pixel were plotted as a heatmap to represent the propagation of calcium signals within the organoid.

### 1.5. Analysis of contractility and beating velocity

A previously published algorithm and MATLAB program (Huebsch et al. 2015) were applied to analyze contractility and beating velocity. Briefly, OCT recordings of heart organoid beating were loaded into the MATLAB program, and the field of view was divided into macroblocks containing 16 by 16 pixels. The quality of the registration of macroblocks at different time frames was calculated by evaluating the mean absolute difference between each adjacent pixel. The motion vectors of each macroblock were generated at different time frames, and the corresponding beating velocities were calculated.

## 3. Results

### 3.1 Longitudinal imaging of hHOs

The OCT setup was used to perform morphological imaging of the hHOs every day from day 1 to day 22 and every other day from day 24 to day 30 (Figure 1B). Figure 2 shows a 3D rendering, *en face* and cross-sectional views, and chamber segmentation of a representative hHO. Small cavities were observed inside 3 out of 32 organoids as early as on day 2. By day 3, after 24 h of Wnt-C59 exposure, cavities were observed in all organoids. OCT allows tracking dynamic development of hHOs throughout the culture period. The human heart is an incredibly complex organ with internal chambers that are paramount to its functionality. The development and growth of these chambers and cavities are of great relevance and interest. In this representative hHO, 36 independent cavities in total were observed inside the organoid on day 14, with volumes ranging from 2.2 × 10^−5^ to 0.059 mm^3^ (Figure 2D). The same organoid was imaged using OCT on day 16 (Figure 2E). The total number of cavities had decreased to 11, number of cavities with small volumes had decreased significantly, and the largest volume of the cavities had increased to 0.086 mm^3^, which indicated active remodeling of the cavities. Segmentation of the chambers revealed complex interconnections of the chambers (Supplementary video 1).

**Figure 2.**
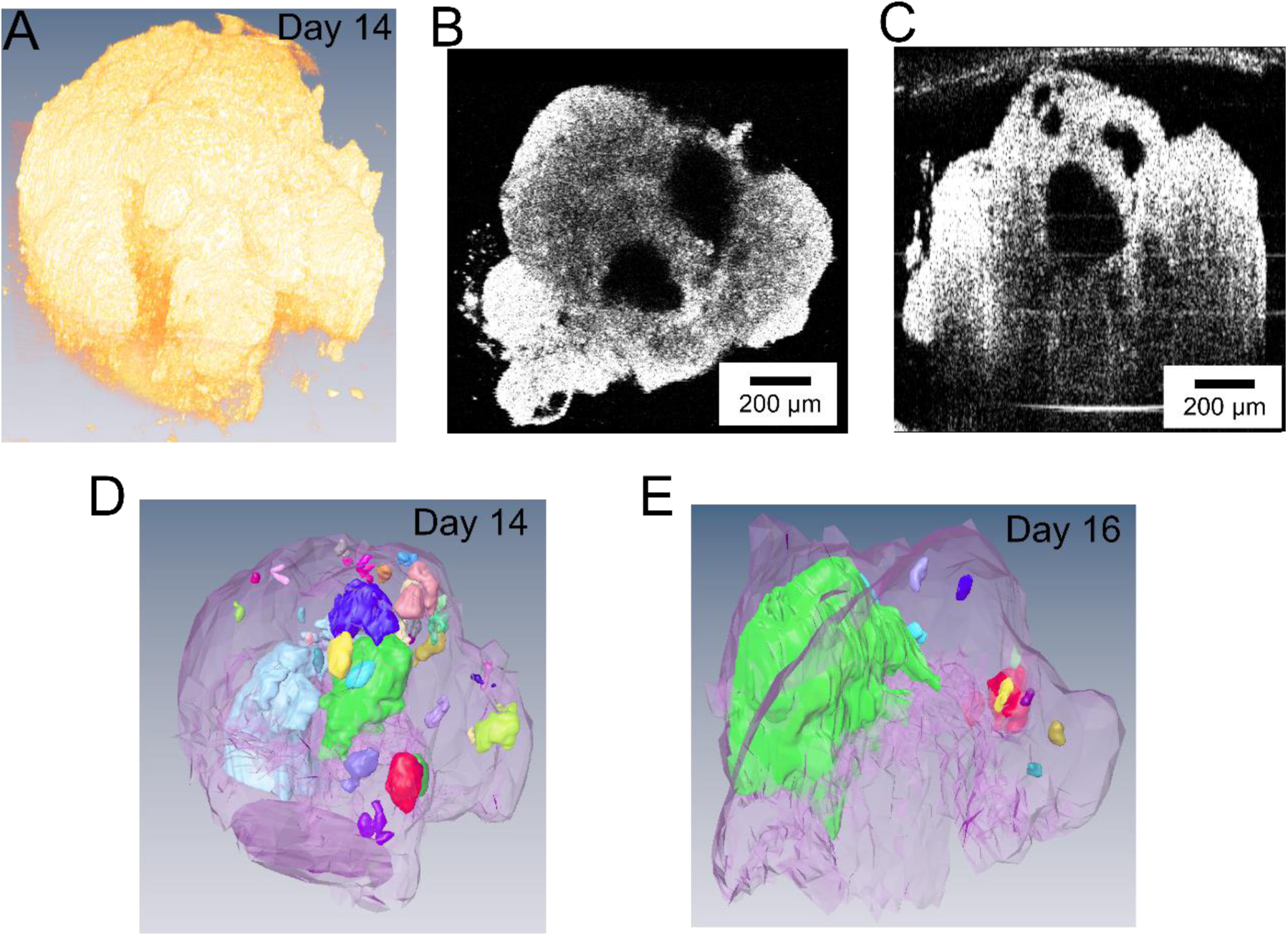
OCT imaging and 3D rendering of a representative human heart organoid. (A) 3D rendering of the organoid. (B) En face and (C) cross-sectional views of the heart organoid. Scale bars: 200 μm. (D) Segmentation of the organoid and the inside cavities. Cavities in different colors are independent (not connected). Panels A-D show the organoid on day 14. (E) Segmentation of the same organoid on day 16.

Label-free and non-invasive nature of OCT imaging allows tracking the dynamic development of the same hHOs on subsequent days. OCT extracts the 3D morphology of a single hHO within minutes, with an imaging depth of around 1 mm, which is not feasible using traditional optical imaging methods. Among all the morphological information obtained from OCT, the development of chambers is of particular interest, as chambers represent crucial structures and functions in human hearts. Figure 3A shows cross-sectional views from OCT imaging of the same hHO. Large chambers within the hHOs are observed starting from day 3-4 in all hHOs. The number, sizes, and overall morphology of the chambers change dramatically. Chamber merging and enlargement are observed from day 6 to day 16. Starting from day 16, both the number and sizes of the chambers decrease. By day 30, no large chamber (>1% of total volume of the hHO) is observed. The sizes of the hHOs (n = 32) were analyzed from OCT scans (Figure 3B). On day 1, hiPSCs formed into embryoid bodies (EBs), measuring 542 ± 57.6 μm (average ± standard deviation) in diameter and 525 ± 59.0 μm in height. After sequential exposure to CHIR99021 (a Wnt pathway activator), BMP4 and Activin A (growth factors), and Wnt-C59 (a Wnt pathway inhibitor), the sizes of the hHOs increased significantly between day -1 to day 4 (p < 0.05, paired t test). The hHOs grew at rates of ∼107 μm per day (linear least squares, R^2^ = 0.961) in diameter and ∼127 μm per day (R^2^ = 0.959) in height from day -1 to day 4 (growth phase I, Figure 3B). Wnt-C59 was removed from the culture medium on day 4 and no significant changes in sizes were observed from day 4 to day 6 (p = 0.38 for height and 0.48 for diameter between day 4 and 5, and 0.30 for height and 0.052 for diameter between day 5 and 6, paired t test). After the culture medium was replaced with RPMI 1640/B27 with insulin on day 6, the hHOs started growing again, but at slower rates of ∼76 μm per day (R^2^ = 0.972) in diameter and ∼60 μm per day (R^2^ = 0.985) in height from day 6 to day 9 (growth phase II, Figure 3B). The hHOs reached a maximum average height of 1327 ± 123.5 μm on day 11 and a maximum average diameter of 1329 ± 180.9 μm on day 9. After the peaks, the sizes decreased slightly and then remained stable.

**Figure 3.**
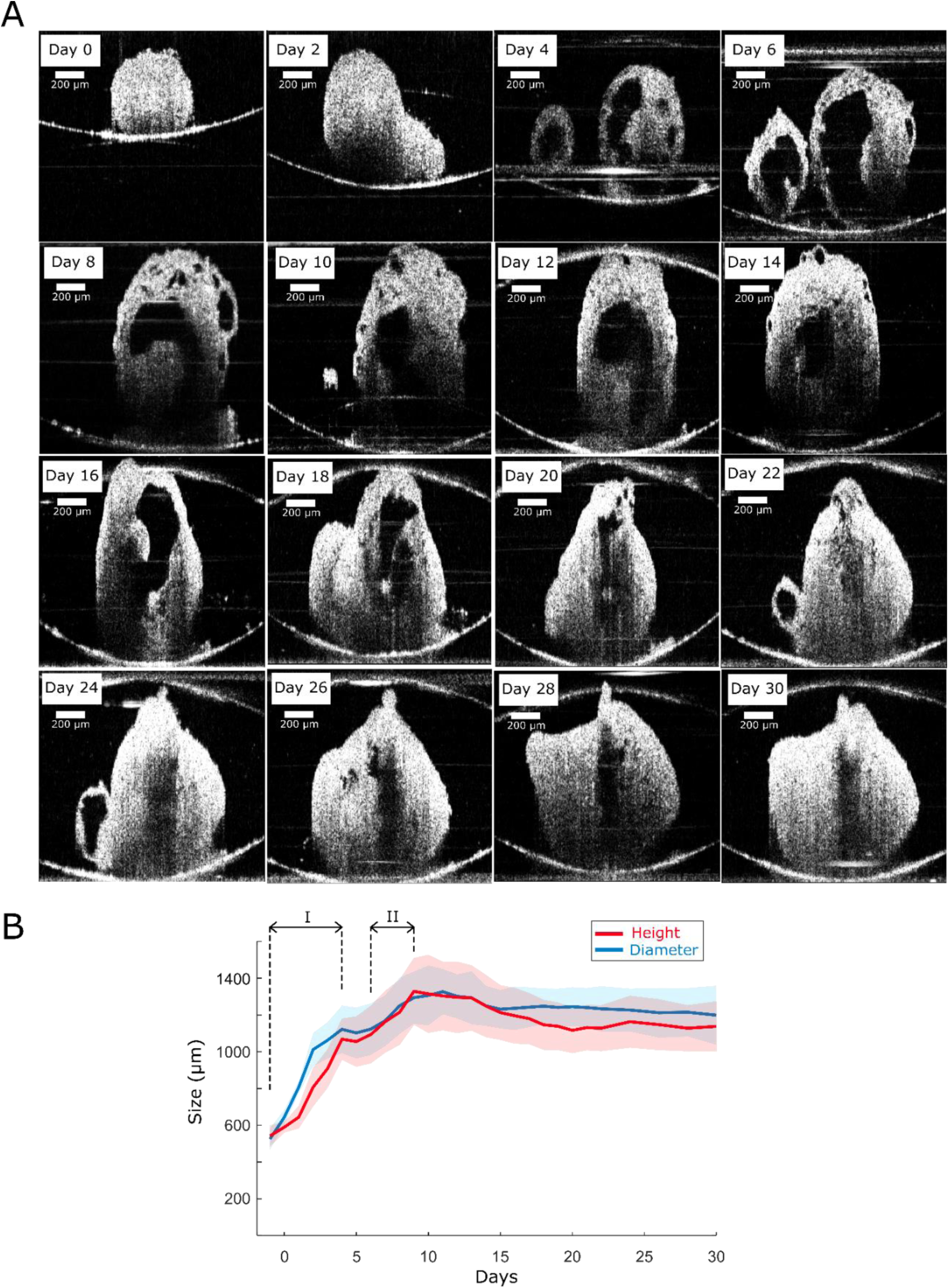
Longitudinal imaging by OCT. (A) Cross sections of the same human heart organoid on different days. Scale bars: 200 μm. (B) Average heights and diameters of human heart organoids on different days. Solid lines represent averages, and shades represent standard deviations. n = 32 organoids. Brackets indicate linear growth phases I and II.

### 3.2 Functional imaging of hHOs

Beating of hHOs was observed as early as a few hours after the culture medium was changed to RPMI 1640/B27 with insulin on day 6. All organoids started beating between day 6 to day 12. Beating of the organoids was recorded using bright field, fluorescent, and OCT imaging (Figures 4 and 5) at room temperature. Spontaneous beating was observed via all three imaging methods (Supplementary videos 2-5). Dynamic changes in fluorescence intensity were observed as early as on day 6, indicating sufficient expression levels of GCaMP to track calcium flux. In our previous work, the percentage of cardiomyocytes was measured as 64.9 ± 5.3% via immunostaining (Lewis-Israeli et al. 2021b). Consequently, variations in fluorescence intensities were not observed in all regions of the hHOs.

**Figure 4.**
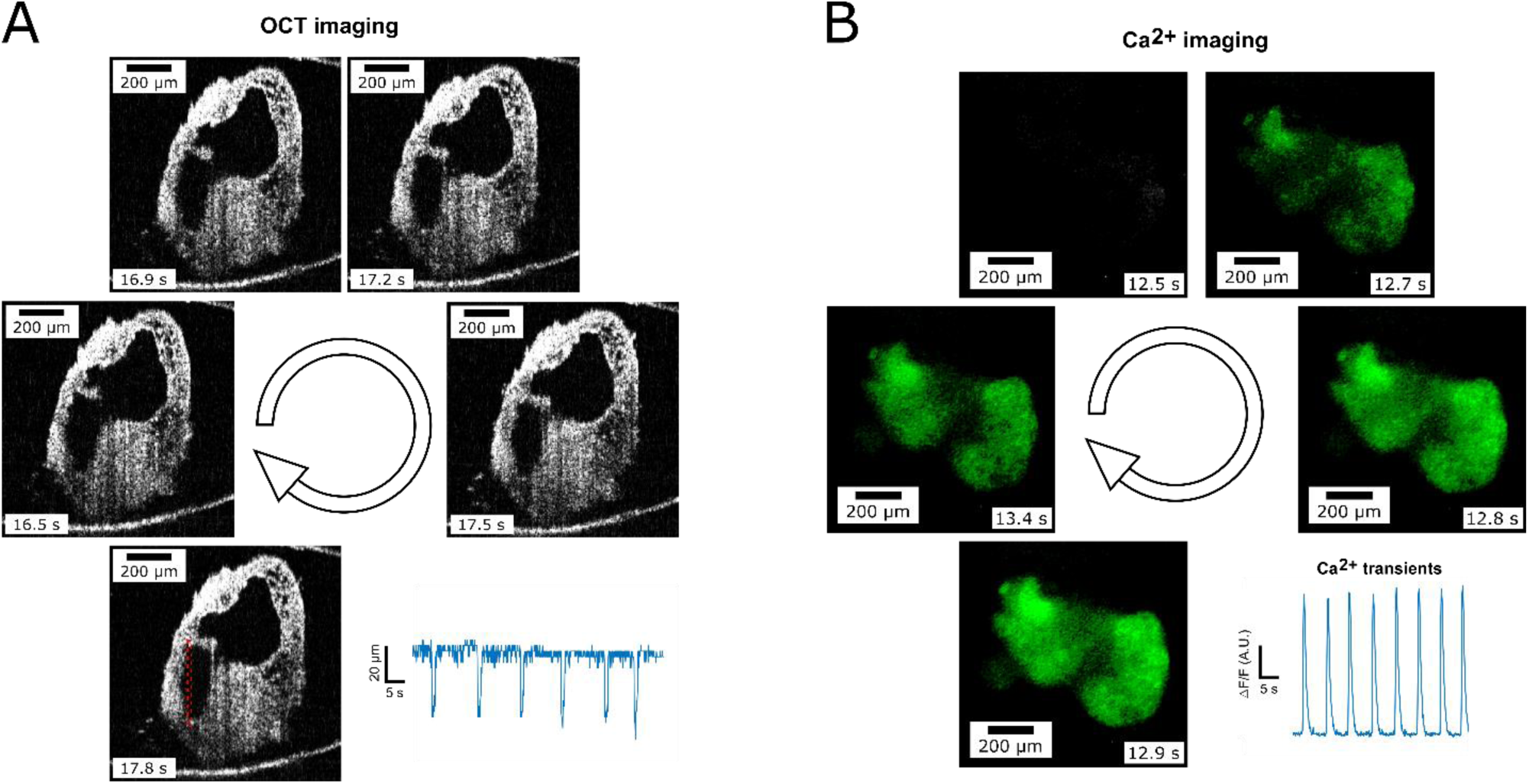
Characterizations of the beating of one representative human heart organoid. (A) hHOs beating characterized by OCT. The blue trace represents changes in the height of the chamber, measured from the red dashed box. (B) hHOs beating characterized by fluorescence imaging of GCaMP6f [5, 6]. The blue trace represents changes in fluorescence intensities of GCaMP6f of the representative binned pixels (see Materials and methods). For panel A and B, 5 representative frames within the same beating cycle were taken from the recordings. All frames were time stamped. The circular arrows indicate the time sequence of each frame group. Scale bars: 200 μm.

**Figure 5.**
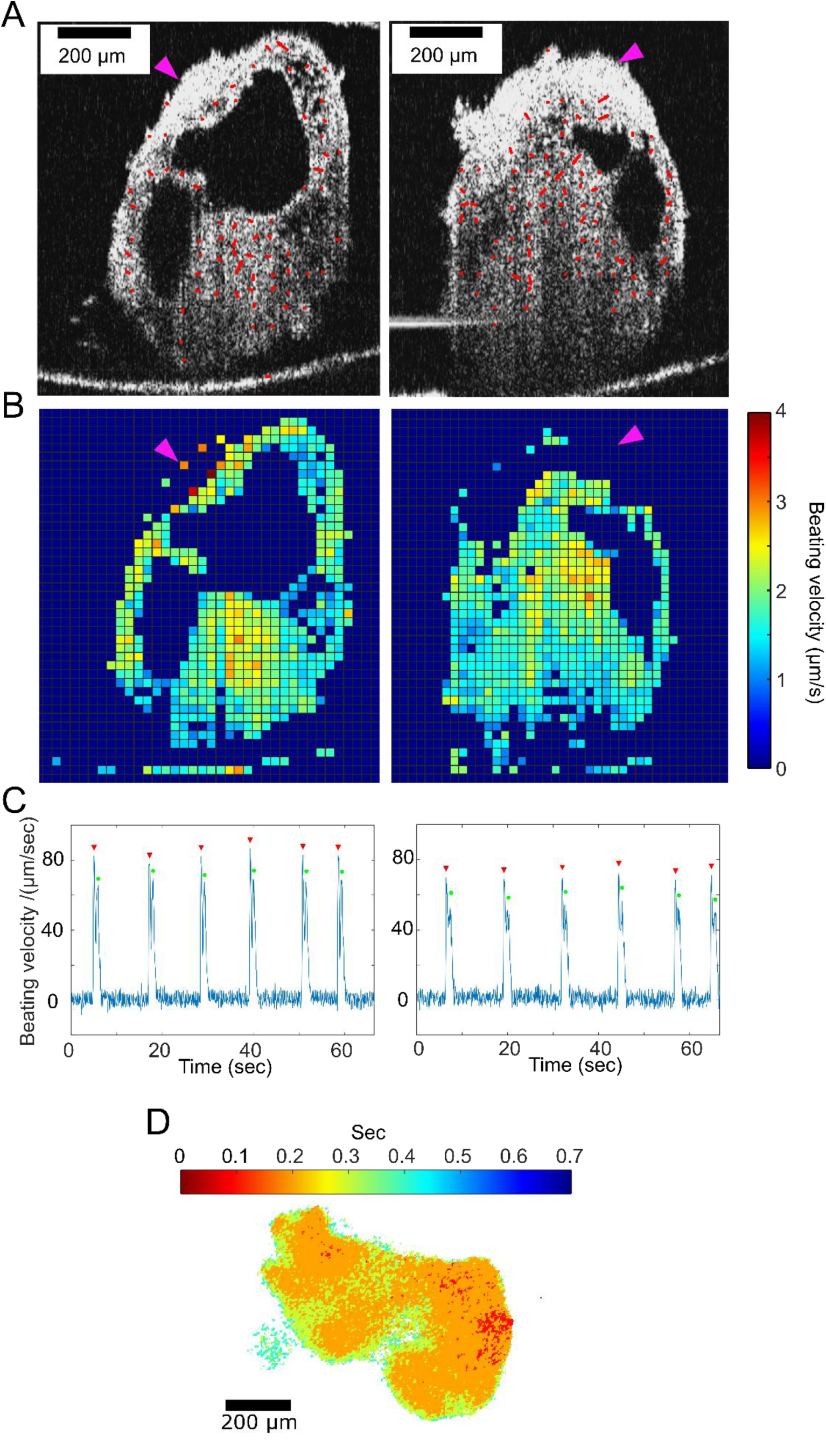
Analysis of beating velocity and calcium signal propagation of one representative hHO. (A) Representative frame of hHO beating recorded by OCT. Red arrows represent motion vectors. The direction and length of each arrow represent the direction and magnitude of the motion vector, respectively. (B) Average absolute beating velocity of the whole recording. Magenta arrows indicate locations with fewer detected motions. (C) Representative traces of beating velocity over time. Contraction and relaxation are marked with red triangles and green circles, respectively. In panels A-C, B-scans on left and right are orthogonal (supplementary video 6). (D) A heatmap depicting initiation of calcium transients of each pixel within the heart organoid. Scale bars: 200 μm.

To track the spontaneous beating of hHOs using OCT, one axis of the galvo system was grounded so that the OCT system acquired repeated B-scan (cross-sectional) images of the hHOs over time (M-mode B-scan imaging). The frame rate of the configuration was 15 fps. Heartbeat rates were calculated by dividing beats within the time series by the acquisition duration. Besides heart rates, size changes of specific chambers during beating were measured using OCT. B-scans containing the chamber of interest were selected for OCT recording, and a box was drawn to include the upper and lower boundaries of the chamber in all frames of the time series (Figure 4A, red dashed box). The A-series of the box was generated to track the movement of the upper and lower boundaries over time, and changes in the height of the chamber were measured by calculating the distance between the two boundaries. Analysis of a representative hHO showed clear chamber contractions during beating (Figure 4A). The average decrease in height of the chamber was 40.9 ± 4.95 μm. The average duration of the contraction was 1.2 ± 0.08 s, and the average beat rate was 11.4 ± 5.22 beats per minute (n = 3 hHOs). Calcium signal propagation was recorded from the same organoid (Figure 4B and Supplementary video 3). The waveform exhibited the typical form of calcium transients, with fast rising and exponential decay (Dittami et al. 2011). The average duration of the calcium peaks was 2.6 ± 0.23 s, and the average beat rate was 15.3 ± 4.93 beats per minute (n = 3 hHOs). The average beat rate was lower than the rate in our previous work (Lewis-Israeli et al. 2021b), which may be due to differences in the stem cell lines or the recording environment.

Another interesting beating pattern observed using OCT involved the valve-like structure seen in Figure 4A. As the lower chamber contracted, the valve-like structure switched to a closed state and physically separated the two adjacent chambers. To confirm that movement of the valve-like structure was not caused by shifting of the B-scan (movement of the hHO in the perpendicular direction of the B-scan), we recorded a time series of the orthogonal B-scan and found the valve-like structure was retained (Supplementary videos 4-6). The valve-like structure inside the hHO was visible due to the 3D imaging capability and high imaging depth of the OCT system. In comparison, standard bright field or fluorescence imaging were not able to resolve the valve-like structure inside the organoid.

OCT recordings of the beating hHOs were analyzed for contractility (Figure 5) using a published algorithm (Huebsch et al. 2015). Frames of the recordings were divided into macroblocks (16 × 16 pixels), and the motion of each macroblock was analyzed (see Materials and methods). Motion vectors in the representative time frames in Figures 5A indicated most macroblocks generated motion during beating of the hHO. Average beating velocities were calculated from OCT imaging. The maximum average beating velocity was observed from the upper portion of the hHO at ∼4.0 μm/s. Motion vectors represented both the directions and magnitudes of the motion of each macroblock. Cellular materials of the hHO exhibited diverse morphology, as observed using OCT. Locations with compact structures (*e*.*g*., densely packed cells) with stronger backscattered signals showed fewer detected motion vectors during beatings (magenta arrows in Figure 5A and B), indicating coupling between the morphology and function of the hHOs. Figure 5C shows representative one-dimensional traces of the beating velocity. Two peaks associated with contraction (red triangles) and relaxation (green circles) of the hHO within each beating cycle are clearly observed. The motion vectors of each macroblock in each frame of the OCT recordings are seen in Supplementary videos 4-5. These OCT recordings reveal beating patterns in the cross-sectional plane. Brightfield and calcium recordings reveal the beating patterns of the hHO in the horizontal plane. The time instants when the calcium signals of each pixel began are plotted as a heatmap in Figure 5D. The earliest calcium signals of the hHO occurred in two spatially separated regions (top left and right red clusters) and propagated from those regions as indicated by the heatmap.

## 4. Discussion

OCT can be dedicated to extract 3D morphological information of organoid models. Because OCT detects intrinsic backscattered light signals, it does not require sample fixation or labelling with contrast agents. The fast, label-free, and non-destructive nature of OCT introduces minimal disturbance to samples. Thus, longitudinal characterization of same samples can be performed via OCT, which yields more faithful observations of morphological development and reduces the number of organoids needed for each experiment. We demonstrated these advantages by monitoring and measuring the development of chambers within hHOs on different days (Figures 2 and 3).

Another interesting feature of the hHOs is their spontaneous beating patterns, which can be characterized via patch clamp, multi-electrode array (MEA), bright field imaging, calcium imaging, and OCT. Each of these methods reveals different information regarding spontaneous beating of the hHOs. Patch clamping focuses on changes in the cellular membrane of individual cardiomyocytes, whereas MEA monitors field potential during beating. Beating velocity, propagation and calcium transients can be extracted via bright field and calcium imaging (Huebsch et al. 2015). Due to limitations of imaging depth, traditional brightfield and calcium imaging can monitor beating of only the top thin layer of the heart organoid. OCT enriches the information by adding chamber contraction and beating patterns in the vertical direction (Figures 4 and 5). In addition, we observed a valve-like structure via OCT. It switched between open and closed states and separated two adjacent chambers during contraction (Figure 4 and supplementary videos 4-6). Another possible way to characterize the function of the valve-like structure is to express fluorescent markers in cells enriched in human heart valves to visualize such structures using live fluorescence imaging. However, this approach is still limited by the imaging depth of fluorescence imaging. It also introduces complexity related to the expression efficiency of the fluorescent marker in the valve-like structure in the hHOs. OCT advantageously visualizes beating patterns in the vertical direction and beyond the imaging depths of brightfield and fluorescence imaging. The transverse and axial resolutions of the OCT we used were ∼7 μm and ∼4.9 μm, respectively. By upgrading the OCT setup with a super continuum laser and a spectrometer with a 2048-pixel camera, we further improved the transverse resolution to ∼3.48 μm and the axial resolution to ∼2.36 μm. High resolution OCT images allow us to precisely characterize the morphology of hHOs, such as cavities of various sizes and shapes, which are related to the development of heart chambers and vascularization, and to monitor beating patterns, including chamber contraction and motion of valve-like structures.

In our previous work (Lewis-Israeli et al. 2021b), self-assembling hHOs were demonstrated to be a reproducible and robust *in-vitro* model of the human fetal heart. The organoid consistently yielded multiple cell types, including cardiomyocytes, epicardial cells, endocardial cells, fibroblasts, and endothelial cells. The combination of the growth factors BMP4 and Activin A with Wnt pathway modulation led to formation of chambers and vascularization within heart organoids. For 3D human heart cultures generated via direct assembly, cardiomyocytes can be differentiated and harvested from monolayer cultures. Thus, the maturation state of the cardiomyocytes can be precisely controlled (Lewis-Israeli et al. 2021a). Our heart organoid model currently replicates the human fetal heart, as revealed by the gene expression profile and fetal-like morphology of the cardiomyocytes. It will be worthwhile to explore maturation protocols and strategies to guide organoids into more mature stages, where morphology and functions recapitulating adult human hearts are expected. The observations that the size and number of chambers within the organoid decrease with time, and the occurrence of a valve-like structure in only some organoids, might be results of inappropriate media conditions to further the growth and maturation of hHOs after a certain stage, a topic inviting further investigation. Organoid models are evaluated from many aspects, and OCT plays an essential role in providing their 3D morphological and functional evaluations. The formation of heart chambers requires complex biological signals, cellular interactions, and different myocardial progenitor cells (Tan and Lewandowski 2020). It involves formation of the heart tube, heart tube looping, and separation into four defined chambers (Buckingham et al. 2005; Später et al. 2014). The heart exhibits distinctive morphology at different stages, and morphological characterization of organoids using OCT is crucial to developing heart organoid models for specific developmental stages. Beyond the formation of functioning chambers, our observations of heart valves, vascularization, and 3D beating patterns provide solid validation of the organoid models. This work further demonstrates OCT’s promising potential for tracking these features of the hHOs.

## 5. Conclusion

OCT shows advantages in longitudinal characterization of the 3D morphology and beating patterns of hHOs. Our heart organoid model currently replicates the human fetal heart, and future work in exploring maturation protocols to model the human heart at subsequent developmental stages is of great interest. The hHOs protocol and OCT both show promising potential in studying the human fetal heart and serve as a starting point for investigating human hearts at different developmental stages.

## 6. Acknowledgements

Work in Dr. Zhou’s laboratory was supported by the National Institutes of Health (NIH) grants R01-EB025209 and R01-HL156265. Work in Dr. Aguirre’s laboratory was supported by NIH grants under award numbers K01HL135464 and R01HL151505, by the American Heart Association under award number 19IPLOI34660342, and by the Spectrum-MSU Foundation. The authors thank Dr. Nathaniel Huebsch and his lab for sharing the WTC hiPSC GCaMP6f cell line and lab equipment, and for valuable discussions.

